# The last battle of Anne of Brittany: solving mass grave through an interdisciplinary approach (paleopathology, anthropobiology, history, multiple isotopes and radiocarbon dating)

**DOI:** 10.1101/2021.02.22.432237

**Authors:** Rozenn Colleter, Clément Bataille, Henri Dabernat, Daniel Pichot, Philippe Hamon, Sylvie Duchesne, Françoise Labaune-Jean, Stéphane Jean, Gaétan Le Cloirec, Stefania Milano, Manuel Trost, Sven Steinbrenner, Marine Marchal, Céline Guilbeau-Frugier, Norbert Telmon, Éric Crubézy, Klervia Jaouen

## Abstract

Mass graves are usually key historical markers with strong incentive for archeological investigations. The identification of individuals buried in mass graves has long benefitted from traditional historical, archaeological, anthropological and paleopathological techniques. The addition of novel methods including genetic, genomic and isotopic geochemistry have renewed interest in solving unidentified mass graves. In this study, we demonstrate that the combined use of these techniques allows the identification of the individuals found in two Breton historical mass graves, where one method alone would not have revealed the importance of this discovery. The skeletons likely belong to soldiers from the two enemy armies who fought during a major event of Breton history: the siege of Rennes in 1491, which ended by the wedding of the Duchess of Brittany with the King of France and signaled the end of the independence of the region. Our study highlights the value of interdisciplinary approaches with a particular emphasis on increasingly accurate isotopic markers. The development of the sulfur isoscape and testing of the triple isotope geographic assignment are detailed in a companion paper [1].

## Introduction

While mass graves are often key historical and sociological markers, with strong incentive for archeological and/or forensic investigations, little historical records usually accompany the hasty burial of corpses. Existing historical records associated with mass burials are often biased: as the Churchillian adage goes, “History is always written by the victors”. In addition, those records usually document the life of the leaders and not that of the physical actors of the battles. The identification of individuals buried in mass graves has therefore long benefitted from archaeological [2–4], anthropological [5–8], genomic [9–11] and isotopic geochemistry analyses [12, 13]. These analyses may specify the context and causes of death (wars, epidemics) but without artefacts in war mass graves, the camps of the buried remain unknown. Isotope ratios of elements incorporated at different life periods in mineralized tissue (bones and teeth) of an individual are likely to document geographical provenance. Here, we demonstrate, for the first time, that the combined use anthropological, pathological, historical and isotopic data, coupled with advanced isotopic models [1,14,15] and existing genetic information [16], can uncover the identity of human remains found in two Breton mass graves while one of those methods alone would have failed to reconstruct their story.

Stable isotope geochemistry has become a popular and effective tool to investigate the diet, childhood or adolescence origin and mobility of buried individuals [17–25]. Nitrogen and carbon isotope compositions (*δ*^15^N and *δ*^13^C, respectively) can reveal key insights into the diet of unknown individuals buried in mass graves, which can be sometimes linked to their social status [26, 27]. Oxygen (O) and strontium (Sr) isotopes have been applied to many archeological studies to determine geographical provenance of recovered human remains or artifacts [12,28–39]. These provenance applications rely on the natural and predictable spatial variability of O isotope composition (*δ*^18^O) and Sr isotope ratio (^87^Sr/^86^Sr) with climate and geological variables, respectively. Advances in ^87^Sr/^86^Sr isoscapes have made it possible to use this isotopic system in combination with *δ*^18^O values in dual isotopes probabilistic geographic assignments [40]. Both isotope systems are generally analyzed in dental enamel, and therefore reveal information on the early life of the individual studied. Sulfur (S) isotopes are another isotopic system that can provide additional independent information to assess the provenance of unknown individuals. The S isotope composition (*δ*^34^S) are known to vary on the landscape with the distance to the coast. We developed in association to the present study an isoscape predicting sulfur isotope composition (*δ*^34^S) across the landscape to use them in multi-isotope probabilistic assignments with oxygen and strontium isotopes [1, 34]. Sulfur isotopes are performed on the collagen contained in the dentine and the bones. In order to have information chronologically closer to that brought by Sr and O isotopes, the triple geographical assignments of our model have been based on dentine values, also revealing information of the childhood and adolescence of the individuals. Bones were also analyzed, and reflect the location of the individuals 10 to 20 years prior to their deaths.

Using an interdisciplinary approach, we investigate the human remains of two undocumented mass graves found in Rennes, Brittany (France). We argue that the exhumed skeletons were soldiers who perished during a major event of Breton history: the Siege of Rennes (1491) representing the last battle for the Duchy independency. This war represents the ultimate battle of the French-Breton war which involved many European forces (England, Spain and the Holy Roman Empire in addition to France and Brittany), and is famous in modern Brittany as it signified the end of the independence of the region. We demonstrate that one of the two graves is linked to the allies of the Duchess Anne of Brittany, whereas the other likely contains remains from members of the French Royal army. Our study sheds new light on this poorly described conflictual event and adds to the historical archive especially on the composition of armies, hitherto unknown. The results underline the power of interdisciplinary approaches to shed new lights on understudied conflicts or events, particularly those using multiple isotope tracers.

## Historical and archaeological contextualization

### Historical background

In the 15th century, the Breton population was estimated at more than one million inhabitants with Nantes (14,000 inhabitants), Rennes (13,000) and Vannes (13,000) being the main cities [41]. At the time, the relative political stability of the Duchy corresponded to a period of prosperity. This stability is illustrated by the construction of many religious buildings, such as the convent of the Jacobins de Rennes [42] and aristocratic manor houses. The city’s walls were also extended between 1421 and 1476 [43]. This prosperity is partly due to the policy of the Montfort family who held the duchy and tried to create a princely state independent of the kingdom. By staying as far away from the Hundred Years’ War as possible, the Dukes promoted peace to a certain extent and economic development based on the development of canvas making and maritime trade.

### The war of Brittany and the siege of Rennes in 1491

The re-establishment of the Kingdom of France after the Hundred Years’ war and the French royal will to impose the liege homage to the Breton Duke triggered a less favorable context for the Duchy of Brittany. The end of the 15th century was thus characterized by a disastrous situation between new war events and their financial and diplomatic consequences as well as new plague episodes. Several reasons led to the conflict against the King of France Louis XI: (i) the restoration of royal power and its willingness to impose itself on Brittany; (ii) divisions within the Breton nobility and (iii) a ducal policy of support for certain revolts against the King of France. Moreover, Duke Francis II, from the Montfort dynasty, had no male heir. According to the treaty of Guérande, at the end of the Breton succession war (1356), the throne should then pass to the Penthièvre dynasty. The new King of France, Charles VIII, son of Louis XI, bought the succession rights to a Penthièvre comtess and claims Brittany whereas the Duke Francis II formalized the position of his daughters as legitimate heiresses with the support of the Estates of Brittany. Tension arose and war broke out in 1487. The Royal Army invaded Upper Brittany, controlled several cities and crushed the Ducal Army at Saint-Aubin-du-Cormier on 28 July 1488 [44, 45]. Then, the Orchard Treaty put an end to the policy of independence of the duchy and stipulated that the two daughters of Duke Francis II (Anne and Isabeau) could not marry without the King’s approval. Nevertheless, other rulers such as Henry VII of England or Ferdinand of Aragon opposed this decision and supported Duchess Anne (**Fig. 1A**). When her father died, Anne ascended to the ducal throne at the age of less than 12 years under difficult conditions. At the end of 1490, she married by proxy - without the authorization of Charles VIII – in order to obtain an ally: Maximilian of Habsburg (**Fig. 1A**). He was the contender for control of the Holy Roman Empire and was at war with the King of France. The latter, opposed to this marriage, brought his army into the duchy. Anne received only limited military assistance from her allies and husband. In addition, the captain of her army, Alain d’Albret, turned his cloak after she wed Maximilian of Hasburg instead of him. In 1491, he used the English soldiers sent by Henry VII to attack the German mercenaries sent by Maximilian of Austria. Anne de Bretagne fought back using the Spanish men sent by Ferdinand of Aragon [46]. A large part of the duchy is occupied by the royal army led by Louis de la Trémoïlle which came to besiege the duchess in 1491, entrenched with the remains of her army in the city of Rennes. The Duchess Anne was then forced to accept a marriage with the King of France by a peace treaty signed at the Jacobins convent of Rennes. From 1532 onwards, the Duchy of Brittany became a French province, entering the royal domain while retaining autonomy with provincial states and then its own Parliament and important privileges, particularly fiscal.

**Figure 1.**
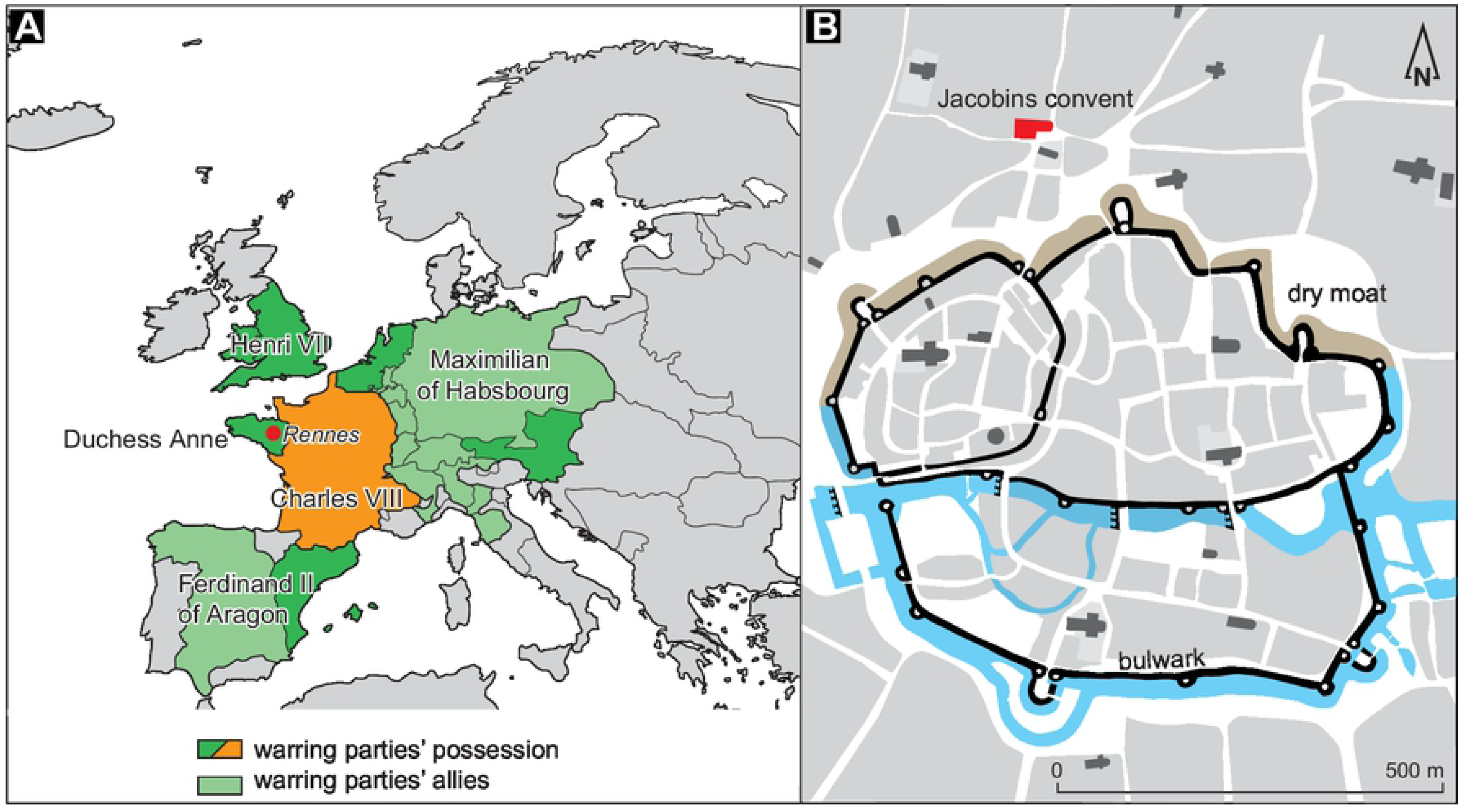
**Location of the site. A**. Situation in Europe in 1491 and the warring parties involved; **B**. Location of the Jacobins convent outside the walls of the city of Rennes.

### The armies involved

Archival sources are patchy on this wartime episode and the studies begun on the state of the forces involved focus more on the 1487-88 campaign and the battle of Saint-Aubin-du-Cormier than on the siege of Rennes. In the 15th century, the Breton army was reorganized by the Dukes. Under Francis II, it initially included a very small number of ducal guards and “Compagnie d’ordonnance” composed of several types of combatants. The whole is based on a heavy cavalry, accompanied by archers, coutiliers and others, forming a small paid army which will develop over the years. In addition, various troops from the feudal service with mismatched combat skills joined the professional army. In 1480, the Duke reformed the Franc-archers (Free Archers), a militia which had been created in 1425 in the Breton parishes. This reorganization aims at improving their fighting skills. Finally, the list of the Breton combatants also includes the mercenaries, who were sent by allies (e.g., Maximilian, Henry the VII, Ferdinand of Aragon) and recruited by the Breton army. The theoretical number of soldiers would be 11,300, but historiographer A. Bouchart suggests there were more than 18,000 men [45]. In any case, the army was decimated in Saint-Aubin (6,000 deaths on the Breton camp). Consequently, the troop around the duchess Anne during the siege of Rennes was very small. The Breton army was supported by the Rennes militia and by a corps of German mercenaries (Landsknecht) sent by Maximilian [47]. English and Spanish mercenaries might also have been sent by Henry the VII and Ferdinand of Aragon [48, 49]. The defense also likely included trained artillery, a crucial resource for the defense of the city.

In contrast to the Breton camp, the royal army commanded by La Trémoïlle was modernized under the leadership of Charles VII and his successors. The heart of the army was formed by the “Compagnie d’ordonnance” of 600 men on horseback whose members were paid, which is an indication of the fighting skills of these individuals. In addition to this company, the King of France called in his Banners. The army also included Franc-archers, mercenaries, and a powerful artillery. The troops involved in the battle of Saint-Aubin du Cormier in 1487 reportedly numbered about 13,500 men [45].

### The Jacobins convent

The Jacobins convent is an emblematic place in the city of Rennes, capital of Brittany, at the western end of Europe (**Fig. 1A**). This community of Dominicans was founded in 1368 by a bourgeois from Rennes but very quickly this foundation was taken over by the Duke Jean IV de Montfort, which undoubtedly contributed to its influence [42, 50]. The convent became a major burial place for the aristocracy of the city but also a local pilgrimage center. Pilgrims used to come to a chapel to pray in front of an icon of the Virgin Mary, “Notre Dame de Bonne Nouvelle” (Our Lady of Good News). The Jacobin convent is located on the west of the Sainte Anne square. At its creation, it was within the parish of Saint-Aubin, outside of the city walls (**Fig. 1B**). The exact configuration of the site during the siege of Rennes remains unknown because, while the church had already been built, the convent buildings (refectory, chapter house, cloister, etc.) were largely renewed in the 17th century [42]. The convent remained under the control of the ducal troops during the siege likely because it was too close to the walls and a city gate (the Foulon’s Gate) to host a royal garrison. Perhaps it also had a strong defense, though archives and archeological findings cannot answer this question. Among the mendicant establishments, the Jacobins convent of Rennes in Brittany appears to be a relatively late foundation (1368) for a major order [50].

## Material

### Sample size

The Jacobins convent was the subject of an entire preventive excavation by the National Institute for Preventive Archeology Research (Inrap) between 2011 to 2013. This excavation of the archaeological site, as well as its study was authorized by order of the prefect (n°35/2011-011). Two burial periods contemporary of the convent (late 14th/16th and 17th/18th centuries) have been identified. Here, the mass graves investigated are exclusively related to the first phase for which 137 individuals were collected (136 skeletons and 1 cardiotaph). Different funeral spaces are individualized (**Fig. 2**). The majority of the tombs from the first phase are located outside of the convent (95/137 or 69%): 64 in the western cemetery and 31 in the cloister garden. Fourteen individuals come from the chapter house, located in the east wing of the convent, 11 from the cloister galleries (3 from the northern and 8 southern galleries), 7 from the church choir, 1 from its adjoining chapel and 8 from the church nave. A lead cardiotaph was also found in a secondary position in the church choir with an inscription dated July 1584 [51]. The funeral recruitment for this phase presents a selection of individuals according to the age of death: 115 adults (of whom 74% are male and 26% female) and 22 children (**SI Text section Physical anthropology**). The individuals between birth and four years of age are under-represented, unlike adults who died between 20 and 59 years of age. The number of men is also too large compared to women for normal mortality (highly significant difference with *p* = 0.0008958; chi-squared test, also written as χ2 test = 11.031).

**Figure 2.**
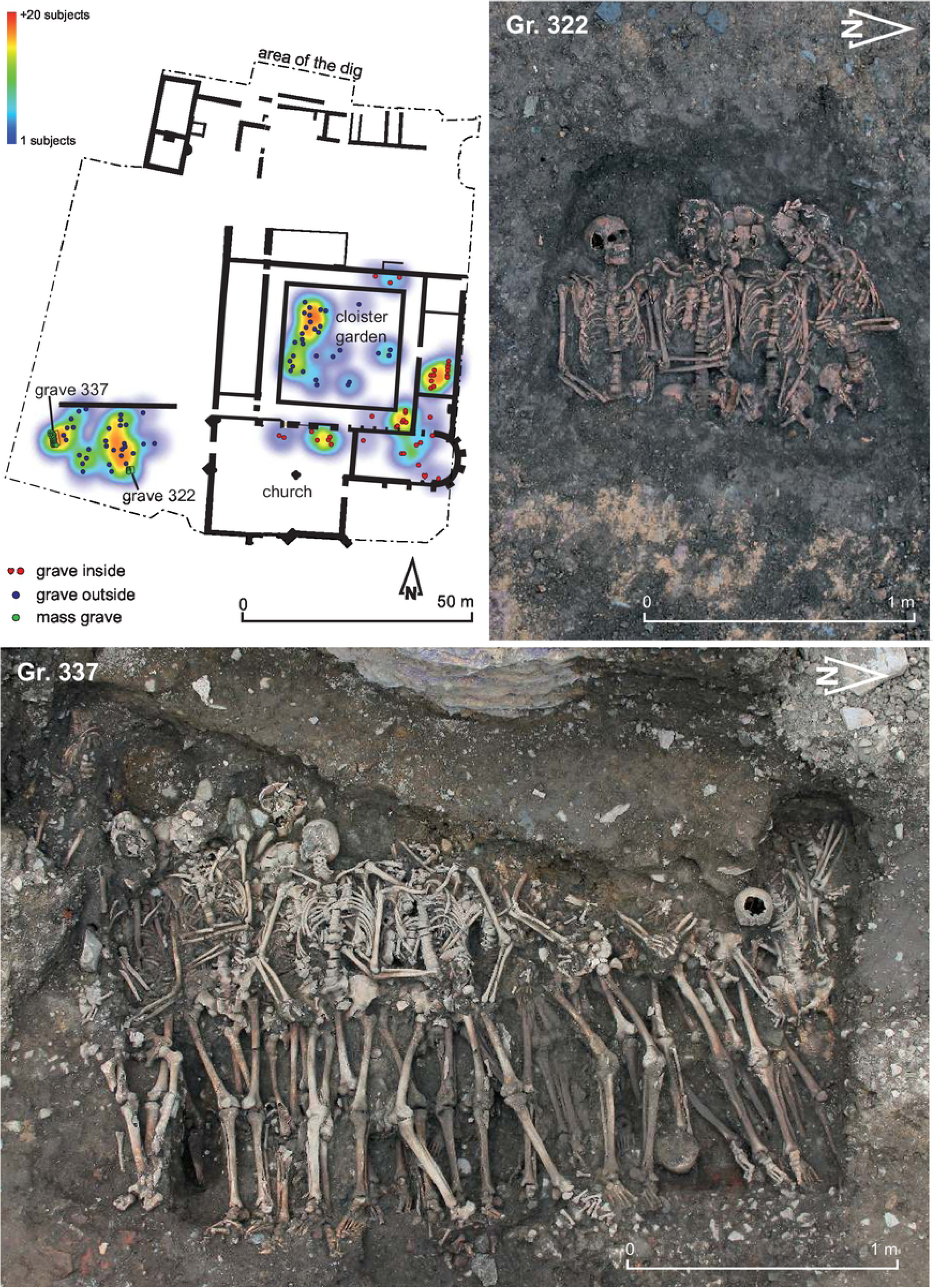
**Distribution of burial**s according to their location (points) and density (heatmap) in the Jacobin convent from the late 14th to the 16th century; photos of archaeological excavation of the mass Gr. 322 and 337.

The two mass graves come from outside the convent (**Fig. 2**). One includes 4 individuals (Gr. 322) and the other at least 28 (Gr. 337). The first tomb was damaged by landscaping work and the construction of a wall, destroying the individuals’ lower limbs. The other pit, a kind of rectangular trench, was only partially dug because of the constraints of the excavation, but at least three-quarters of which was excavated based on additional selected drilling. Gr. 337 was filled with sediment including many terracotta elements, shale blocks and some fauna bones after the funeral depot. The filling partially destroyed parts of an old street in the Roman city. Later, the tomb was disturbed by the installation of a wellbore in its western part, intersecting the upper part of many skeletons. The sepulchral pit was excavated over a length of 2.60 m, with a maximum extension, after drilling, estimated at 3 m. Twenty-eight individuals in primary position are identified but the overall estimate is 32 or even 35 individuals.

### Sampling strategy, buried individuals

The location of the tomb within the convent buildings served as a basis for assessing social groups of the deceased (cemetery vs. church for the French historian Michel Vovelle [52]). Indeed, it is the proximity to certain architectural elements that is important in Christian burial tradition [53, 54]. Individuals buried inside the convent-choir and nave of the convent church and chapels-are considered to have a privileged status. According to historical sources [55], those coming from the Chapter house would probably be clergymen (Dominican friars). All these individuals are grouped in Group **I** (Inside) and represent 31% of the total number of individuals studied (42/137). Forty-two individuals (41 skeletons and 1 cardiotaph) were counted. A second group, less favored than Group **I,** comes exclusively from the surroundings of the convent. These individuals are grouped in Group **O** (outside) which represents 46% of the numbers of individuals studied (63/137). The two mass graves, found outside the convent, are individualized in the study in a “**Mass Grave**” group.

In the absence of dated artefacts, we sent two samples for radiocarbon dating through Accelerator Mass Spectrometry using a selection of human remains from each tomb. We used a fragment of forearm from subject 20183 (Gr. 322) and from the clavicle of 20769 (Gr. 337). The samples have been analyzed by the company Beta Analytical, Floride, USA. The collagen was extracted with alkali by the company in 2015-2016 (**SI Text section Datation**).

For the isotopic analysis, we selected the bones of the individuals of the sepulture Gr. 322 (4/4) and Gr. 337 (22/28), and one tooth for each individual with cranial remains (4 teeth for the Gr. 322, 6 teeth for the Gr. 337). All the teeth were sampled for O and Sr isotope analyses, and the tooth root was also sampled for C, N, S isotope analyses. Oxygen, strontium and sulfur isotope analyses are described in our associated article [1]. The associated local fauna recovered in the grave and previously studied by Jaouen et al. [26] were also sampled. We also added two local individuals (Louise de Quengo and Louis du Plessis from the 17th c. [56, 57]), since no oxygen isotope analyses were previously performed on humans for this site.

## Methods

### Physical anthropology and funeral practices methods

The sex determination was made from observations [58] and measurements made in the pelvis (i.e., Probabilistic Sex Diagnosis in French [59]). Age at death was estimated for adults from observations of the sacropelvic surface [60] and that of adolescents and sub-adults from the stages of tooth eruption [61, 62] and bone maturation [63, 64]. The statures were calculated according to Ruff’s equations [65] from measurements of the maximum lengths of the femurs and humerus. They follow a normal Law (Shapiro-Wilk test, W = 0.9910). An ANOVA and Tukey tests were calculated to determine significant relationships between groups.

The determination of funeral practices is based on archeo-thanatological observations in the field [66]. The preservation or not of all anatomical connections, the observation of any preserved volumes, the unstable balances or linear effects on human skeletons, all provide information about the burial method and funeral architecture.

### Lesions and paleopathological methods

The paleopathological study was carried out on the basis of a classical macroscopic bone examination through the identification of lesions and their possible interpretations. The list of lesions includes trauma, specific and non-specific infections, non-specific stress markers, congenital anomalies and malformations, osteoarthritis lesions and other inflammatory or degenerative diseases, as well as entheses’ changes (i.e., evidences which cannot be linked to a specific activity) [67–72]. In the case of multiple burials, the location of the trauma was compared with that of the organs, serous and soft tissues (arteries, veins, peripheral nerves, muscles, tendons, ligaments, etc.) in order to assess the consequences of the injuries.

Some of the identified bone lesions were observed under an epifluorescence macroscope using the protocol of Capuani et al. [73] The epifluorescence macroscopy combines the advantages of macroscopy (large workspaces, large working distances and accurate reproduction) with high-resolution fluorescence technology. The function of the epifluorescence macroscope is to irradiate the sample with a desired range of wavelengths and then separate the fluorescent light emitted from the excitation light. Incident light is produced by the passage of source light through an excitation filter, which selects the wavelengths of interest. The wavelengths transmitted by the excitation filter are deflected by a dichroic mirror and pass through the lens to the sample, which is therefore illuminated from above. The light emitted by fluorescence is collected by the lens. It then passes through the dichroic mirror and an emission filter that blocks the excitation wavelengths reflected by the sample, and reaches the eye or camera. The excitation filter, dichroic mirror and emission filter are assembled in a cube. Several interchangeable cubes are arranged in a rotating turret, the filters and mirror being adapted to the fluorescent body to be analyzed.

The description of the wound marks was standardized to facilitate their interpretation: a stab wound with a knife consists of i) two walls (parts of the bone in contact with the blade); ii) two banks (meeting lines between the wall and healthy tissue); iii) two profiles (cutting planes of the lesional groove) and iv) a bottom (deepest area of the notch) (**SI Text section Lesions and paleopathological data**; fig. S6). Bone debris resulting from the passing of a blade and the destruction of the bone might be present in different areas of the notch. A “*rattail*” appearance indicating where the blade left the bone is an indication of the end of a lesion.

### Isotope analyses

Part of the isotope analyses have been performed in two previous publications on the Jacobin convent [26, 74] as well as the associated publication [1] for the local fauna and humans. For C and N isotopes, 7/26 individuals were first published in Colleter et al. [74], and we here analyze the rest (20/26) of the individuals. Sr isotope ratios of teeth were previously published for 3/10 individuals, and we here analyze the 6 other individuals. All the oxygen isotope data (10/10) on the teeth are from this study.

Tooth and bone cleaning and sampling for C, N, S isotope analyses of the collagen were performed using the protocol previously described in Colleter et al. [26, 74]. The collagen was extracted using the protocol of Talamo and Richards [75] and the C/N isotopes were analyzed using a Delta 5 EA-IRMS. Standards were analyzed along with the samples and they showed expected ratios (table S9).

Sulfur, oxygen and strontium isotope analyses were conducted in the frame of our associated publication for the local individuals [1]. For the individuals of the two mass graves, we conducted the exact same analyses and methods. Among the 28 individuals of this Gr. 337, we conducted isotope analyses for 6 teeth and 22 bones belonging to 23 different individuals. For sulfur isotopes, 16/22 of the human bone data and 3/6 human tooth data are from this study, and the rest was published in Colleter et al. [74]. For the Gr. 322 and 337, 7/10 Sr isotope analyses were already published in Jaouen et al. [26]. The 3/10 additional Sr isotope data are from this study. All oxygen and carbon enamel isotope data (10/10) are from this study. Eight mg of the remaining collagen extracted for C and N isotope analyses was sent to Isoanalytical for S isotope analyses by EA-

IRMS. Dental enamel was first mechanically cleaned prior to sampling for Sr, O and C isotope analyses. The strontium was extracted using a modified protocol from Deniel and Pin [76] in the clean lab facility of the Max Planck Institute for Evolutionary Anthropology. They were then analyzed at the Max Planck Institute and the University of Calgary using the Neptune MC-ICP-MS. For O and C isotope analyses of the tooth enamel, enamel powder (8mg) was sampled using a diamond drill for each tooth and sent to Iso-analytical LTd (UK) where oxygen and carbon isotopes analyses have been performed by CF-IRMS.

In our associated publication [1], we developed a sulfur isoscape, allowing for the first time to combine these three different isotope systems into probabilistic geographic assignments. We applied this model to the individuals of our two mass graves to establish their geographical origin. Details of the methods can be found in the above-mentioned article.

## Results

### Anthropological and paleopathological results

At the Jacobins’ convent, individuals were buried either within the walls (Group I, for inside) or in the yards and cloister (Group O, for outside). The group I included 8 individuals under 20 years of age and 34 adults (as many men as women when sex is determined) and the group O counted 52 adults (71% males and 29% females with a determined sex) and 11 individuals under 20 years of age (table 1). The Group I presents distortions compared to archaic mortality: (i) an over-representation of adults who died between 30 and 59 years of age and an under-represented of children under 4 years (fig. S1)

Among all the burials excavated contemporary of the convent, only two graves include more than two skeletons (**Fig. 2**). Twenty-nine adults and three individuals between 15 and 20 years old, all men, were counted in these two graves (table S1). The demographic profile clearly shows a significant selection of the buried persons (**SI Text section Physical anthropology**; fig. S1). The first grave (Gr. 322) is located in the middle of the outer western cemetery and contains four individuals buried simultaneously based on the skeleton’s position in the pit (Fig. 2) [77, 78]. The second one appears to be a kind of trench dug at the occidental part of the same funerary space and contains at least 28 individuals (Gr. 337). The first one does not necessarily constitute a mass grave *stricto sensu* since it only contains four individuals [79–81]. Nevertheless, the contemporaneity of the deposits clearly illustrates a brutal and morbid event [78].

Simultaneous to the particular recruitment of the grave (only men over 15), the existence of traumatic injuries observed on the skeletons argues for a soldiers’ burial. The individuals are characterized by numerous perimortem lesions by sharp or spiked objects (**Fig. 3; SI Text section Lesions and paleopathology data**; figs. S7 and S8). Only the skeletons of the mass graves have unhealed lesions and wounds in the upper part of the body (skulls and upper limbs). Trauma sequelae are present on the skeletons of 2 individuals from the Gr. 322 (skeletons 20183 and 20188) and 14 from mass Gr. 337 (fig. S8, S10 and table S4). For this last tomb, it should be noted that the best represented skeletons from the southern part of the pit almost all have lesions. Five skeletons of the Gr. 337 have healed traumatic lesions (fig. S9) without consequence on the death of individuals 11 individuals have trauma from spiked or sharp objects, with more or less secondary lethal consequences. Among those with lethal lesions, two individuals (20764 and 20800) also show healed lesions that suggest a return to combat. For each individual in the Gr. 337 and Gr. 322, we investigated what type of lesions can be observed on their bones, as well as their location which can help to identify in what type of fight they were inflected (riders, backstabbing, etc…). Detailed results are described in **SI Text section Lesions and paleopathology**. Seventeen unhealed blows which reached the bone are identified for the 28 individuals of the Gr. 337 (**Fig. 3**). Some individuals, especially older ones, combine lethal injuries and healed ones (figs. S8, S9, S10 and table S4), which suggests that they regularly fought throughout their life [82, 83]. This observation suggests that those individuals were professional soldiers or mercenaries. Observation of lesions by epifluorescence macroscopic suggests that the weapons causing the injuries were sabers or halberd (**SI Text section Lesions and paleopathology**). In the largest tomb (Gr. 337), the blows that impacted the lower limbs (ankles or thigh) were given from the bottom to the top, which evokes the will to dismount the horse riders (4/17) (table S4). On the other hand, the blows received on the backs of the victims (3/17) and their forearms (5/17) as well as the lesions on the skulls (5/17) suggest the killing of soldiers once on the ground. We argue that the characteristics of the wounds and locations of the blows (head, face and upper limbs) are likely the results of a close combat with knives between two professional armies. Skull blows are often more severe in cases of anti-personal violence and abuse, possibly because the head is considered the seat of identity [84–86].

**Figure 3.**
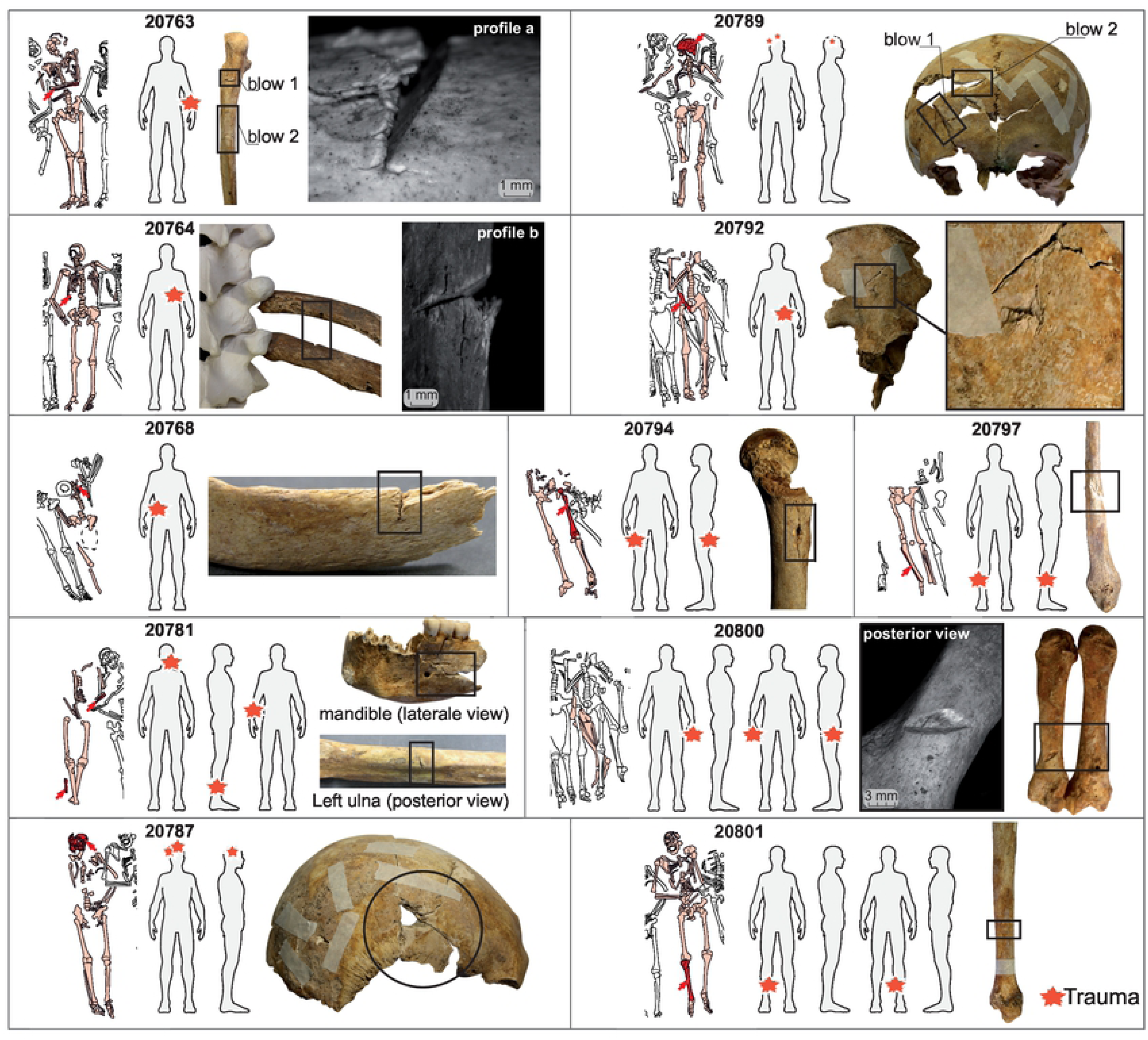
**Lesions and Paleopathological data for individuals recovered from Gr. 337.** Each box represents one individual characterized by its numeric ID (fig. S8). The skeleton schematics represent the reconstructed in situ grave position. Red stars on gray body shape point to the location of traumatic injuries. Black circles and rectangles indicate macroscopic observations of traumatic injuries on bones and selected epifluorescence macroscope photos (profile a, b and posterior view, fig. S6).

The statures of the individuals from the mass graves are higher than the other individuals buried in the convent (1.72 m for burial Gr. 322, 1.68 m for Gr. 337 and 1,61 m for the other; fig. S2A; *p* = 0.00192, ANOVA, Shapiro-Wilk normality test W = 0.99108, *p* = 0.93) (**SI Text section Physical anthropology**; fig. S2A and table S2). Those from Gr. 322 are significantly taller than those from groups **I** and **O** (fig. S2B); the individuals of Gr. 337 are also statistically taller than group **I**. The height standard deviation is also lower in those graves relative to other groups, which shows a certain homogeneity in morphology. If the morphology of individuals reflects both genetic and mesological parameters [87–91], the clear difference between these graves and the rest of the convent suggest a specific recruitment of the buried individuals. In archeological assemblages that are not biased regarding recruitment, average stature can be an indicator of people’s living conditions and their evolution [91–93]. Classically, stature is linked to the intertwined effects of genetics, culture and environmental factors-including diet. Here, on the contrary, the elevated height could reflect a selection of individuals recruited in the army [55].

### Radiocarbon chronology: an event of the 15th century

Assuming that these graves belonged to soldiers who died during combat, we still have to establish the time of the war event. Radiocarbon analyses offer a rather wide range for both tombs, from the middle of the 15th century to the end of the 16th century (**SI Text section datation**; fig. S5). The lack of dating precision is due to an unfortunate plateau effect in the isotopic calibration curve between the 15th and 17th centuries. During this period, Upper Brittany experienced only two conflicts within its borders: the War of Brittany (1487-1491) and the League War (1589-1598). The absence of gunshot injuries, the high number of stab wounds and the absence of historical records argues for a late 15th century conflict [7, 83]. Rennes has only known one major violent episode likely to have caused casualties during this time period. It is dated from 1491AD and it corresponds to the conquest of the city of Rennes by the French forces, all the rest of the Duchy of Brittany being already under their control. The few objects (3 different sets of pearls from the same jet rosary and the lower half of a bell) found in direct contact with the bodies suggest the absence of a complete skinning (careful search and/or theft of corpses and presence of clothing) (fig. S4) [82, 94]. The presence of a rosary probably could illustrate a benevolent attention at the time of the inhumation of these victims [83].

### Carbon and nitrogen stable isotope results

All the isotope results for samples and standards are available in the Supplementary Tables S6 to S9.

The values for the 4 individuals of the mass grave 322 are quite heterogeneous for nitrogen isotopes, ranging from 9.5 (individual 20185) to 13.0‰ (20193) in bones and from 9.7 (20185) to 12.8‰ (20183). Those values are similar in the bones and the teeth belonging to the same individual, except for the individual 20188 which exhibits a higher δ^15^N in bone relative to his M3 (**Fig. 4**). Carbon isotopes values are ranging from −18.7 to 20.1‰ in bones and −19.1 to 20.2‰ in teeth. Nitrogen isotope ratios of the individuals 20183 and 20193 overlap with that of the individuals buried in the church of the nearby convent, which are well-off individuals (**Fig. 4** [74]). Conversely, the individual 20185 exhibit the lowest δ^15^N value of the whole convent, grave 337 excluded.

**Figure 4.**
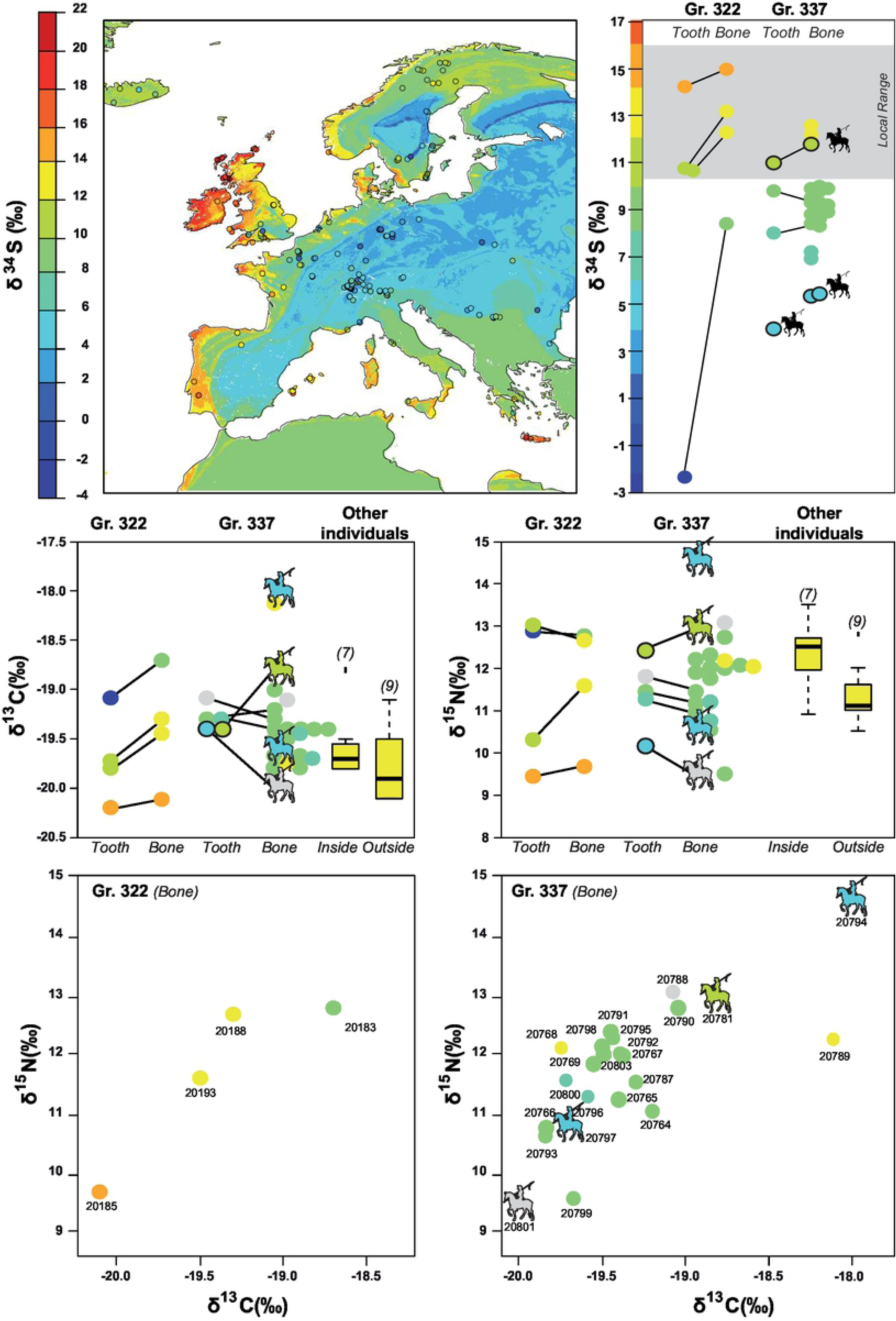
**Carbon, Nitrogen and Sulfur isotopes values within Jacobins’ mass graves. To the top**: Sulfur isotope composition of human remains across Europe according to Bataille et al., this issue [1] and δ^34^S values for individuals of the Jacobins’ mass graves. The first point represents the average δ^34^S values from childhood and is based on teeth roots’ collagen, the second point represents the average δ^34^S values from adulthood and is based on bones’ collagen, the local range represents the animals range according Colleter et al. [74]. **Middle**: Nitrogen and carbon isotope values of the teeth and bones of the individuals from the mass graves 322 and 337, compared to the individuals buried within the convent (inside and outside. **Bottom:** Carbon and nitrogen ratios in the collagen of the bone for the individuals of the mass Gr. 322 and 337. The colors indicate the provenance of the individuals, based on their δ^34^S, grey when values are unknown.

It is indeed in the grave 337 that the lowest δ^15^N values can be found in the bones of the individuals 20799 and 20801. This individual 20801 is one of the four riders identified in the two graves (**Fig. 5**). This second grave also exhibits heterogeneous δ^15^N in the skeletons of the different individuals, with the higher value of 14.6‰ for the individual 20794 who also has been identified as a rider (**Fig. 5**). The values in the teeth are also homogenous in the teeth and bones of the same individuals (**Fig. 4**).

**Figure 5.**
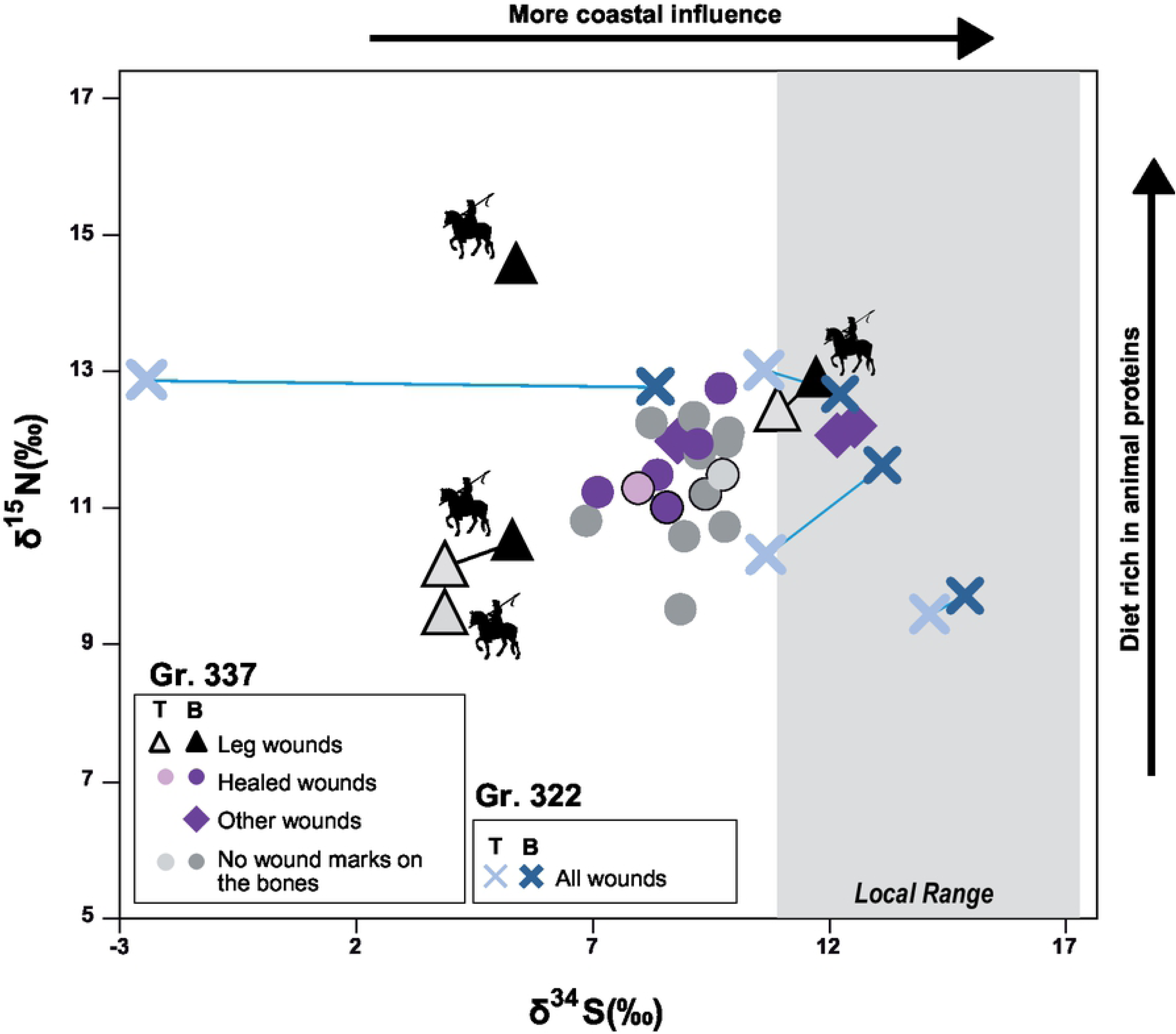
**Nitrogen and sulfur isotope data of the soldiers in the graves 337 and 322** in relation to their wound marks on bones. T stands for teeth values and B for bones.

### Sulfur, oxygen and strontium isotopes

We used *δ*^34^S, *δ*^18^O and ^87^Sr/^86^Sr from bones and teeth of individuals from the mass grave to reconstruct the mobility, childhood and/or adolescence origin and last years of life of selected individuals from each grave. We analyzed *δ*^18^O and ^87^Sr/^86^Sr in dental enamel as well as *δ*^34^S in collagen of teeth and bones for a selection of individuals (tables S6 to S8). Teeth form during a person’s childhood or adolescence and preserve the isotopic composition of that period [95] whereas bones record approximately the integrated isotopic value of the last 10 to 20 years of life [96]. When sampling the tooth root for sulfur while we sampled the crown for Sr and O isotopes, we are sampling two tissues successively forming. The following geographical assignments that we are making therefore relies on the assumption that the individual stayed in the same region at the time of his tooth formation.

#### Grave Gr. 322

Among the 4 individuals of the Gr. 322, 3 have ^87^Sr/^86^Sr, δ^18^O and δ^34^S ranges compatible with local ranges defined in Colleter et al. and Jaouen et al. [26, 74]. Two of the individuals (20188 and 20193) have a tooth δ^34^S value slightly out of the range defined for Rennes area [74] but the δ^34^S values of their bones fit with the local range. This observation suggests that those individuals moved during their life, possibly coming from the more inland region of Brittany on the Eastern part of the Armorican Massif (**Fig. 4**). This moving possibly happened shortly after their adolescence, when they were age of 16-35 years old for the first one and 16-25 years old for the second one as they both died quite young and that the teeth analyzed were M3 (**SI Text section Isotopes data**; table S7). This hypothesis is plausible for the individual 20193 according to its dual ^87^Sr/^86^Sr – δ^18^O isotope assignment (**Fig. 4** [1]). Individual 20188 could i) come from central Brittany ii) come from a totally different region showing similar ^87^Sr/^86^Sr - δ^18^O values as Brittany including the Aragon Kingdom (allied to Brittany during the French Breton war) or the Aquitaine area (part of the French Kingdom, the opponent of Brittany) or iii) be local but have eaten less fish which tends to lower δ^34^S values [97–99]. This latter hypothesis is also supported by the elevated δ^15^N value of the tooth of this individual, which is the highest of the four soldiers in Gr. 322. One soldier (20183) has ^87^Sr/^86^Sr and δ^18^O values compatible with the local isotope ranges but the lowest δ^34^S values in teeth for the two mass graves. This very inland δ^34^S signature is incompatible with the hypothesis of a childhood spent in Brittany. Similarly, the bone δ ^34^S value for this individual is also out of the Brittany range, though it is closer to it (**Fig. 4**). This means that the individual moved during the course of his life, and possibly returned to Brittany within the last few years before his death. As bones turnover slowly over a period of several years, this would explain the intermediate δ^34^S value observed in the bones of this individual. Possible childhood locations in Europe for this individual are: Marnes region (France) or inland Spain. Based on the triple isotope assignment, the most likely place of childhood would be the region stretching from the region of Coulommiers/Melun (South-East from Paris, France), but the lack of additional data prevents us from drawing further conclusions. All the isotopic values for individual 20185 are compatible with the local range, and triple isotope assignments show strong probability of childhood origin in Brittany. Nevertheless, the isotopic values are also compatible with international regions: Northern Spanish coast, Aquitaine coast, Cornwall and Wales. However, since the isotope values for nitrogen, carbon and sulfur are extremely similar between teeth and bones, it is likely that the individual did not move during his life and was local.

#### Mass grave Gr. 337

Among the 28 individuals of the Gr. 337, we conducted isotope analyses for 6 teeth and 22 bones belonging to 23 different individuals. For sulfur isotopes, 16/22 of the human bone data and 3/6 human tooth data are from this study, and the rest was published in Colleter et al. [74]. All δ^18^O and δ^13^C of tooth enamel are from this study. Finally, 3/6 ^87^Sr/^86^Sr human isotope data are from this study, the three other individuals were analyzed in Jaouen et al. [26].

The δ^34^S values in both teeth and bones are lower than the local range. A Nemenyi test following a Kruskal Wallis test shows that the individuals from the Gr. 337 have δ^34^S values significantly different from the individuals buried in the church who were likely locals (group **I**; *p* = 0.0002). Only three of the 21 individuals analyzed from bones (14%) of the mass Gr. 337 have isotopic values compatible with the local range (20781, 20789 and 20768) (**Fig. 4**). One of these individuals (20781) had a tooth preserved and was analyzed for S, O and Sr isotopes. This individual is not compatible with the local range but is rather compatible with western France (Poitou, Charentes area) but also England, the Netherlands and Asturia. For the two other individuals, no teeth were preserved and we could not further assess their region of childhood origin. Consequently, we can only conclude that they lived in a coastal region but not necessarily Brittany (**Fig. 4**).

Based on δ^34^S geographic assignment from bones, individuals from this mass grave have a broad range of origin. δ^34^S values range between 5‰ and 13‰ which span the baseline values of most of the French Kingdom (**Fig. 4**). For the six individuals for whom multiple isotope analysis was possible (**Fig. 6**), all these individuals have ranges compatible with a French origin (6/6). While less likely, these individuals could also come from different foreign kingdoms: 2/6 could come from Brittany (individuals 20787, 20788-although no S isotope values are available for those individuals), half of them could come from England (individuals 20764, 20781 and 20788, either 3/6), all of them from current Spain, but two of them likely come from region that did not belong to Castille and Aragon in 1491 (20765, 20781,), and 5/6 from the Holy German Empire (20764, 20765, 20781, 20788, 20801) (**Fig. 6**). Within the Holy German Empire, two of them are likely to come from the Netherlands and/or Belgium whereas no historical sources identify the recruiting of mercenaries from this region. If from a single political entity, the most likely candidate of origin for the soldiers of the grave 337 would therefore be the French Kingdom.

**Figure 6:**
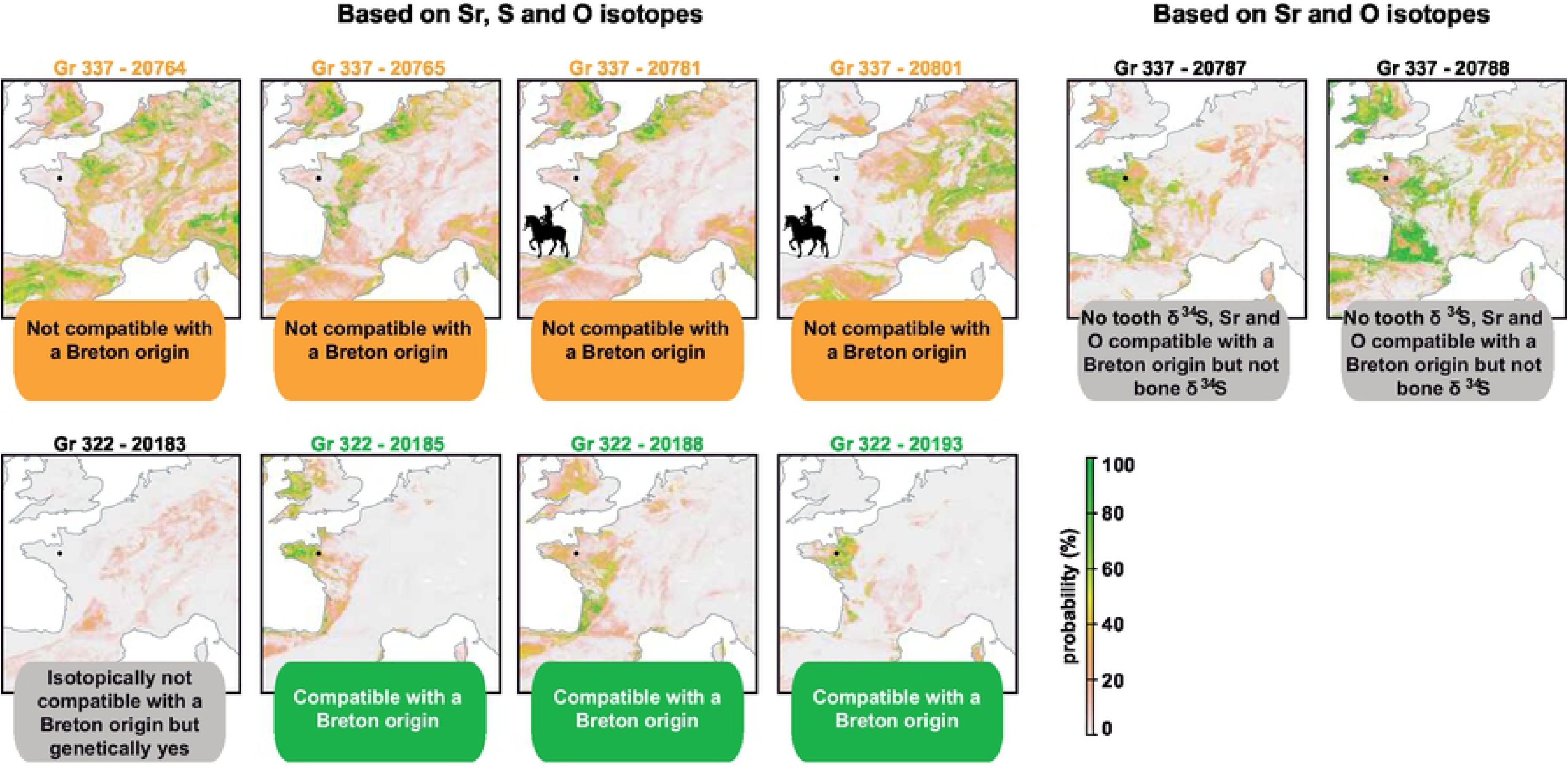
**Maps showing the probability of childhood origin of all individuals** for which teeth were available in the mass graves. Maps of possible geographical origins are calculated by comparing predicted isotope variations on the landscape with isotope analyses from tooth enamel for Sr and O isotopes and from tooth’s collagen for S isotopes [1]. Single, dual and triple isotope geographic assignments were performed for each individual but only the results from triple isotope geographic assignments are displayed. For individuals 20787 and 20788, not enough collagen was preserved for analyzing δ^34^S values and only dual Sr-O isotope geographic assignments are displayed.

The δ^15^N and δ^34^S values of bones and teeth of this group are highly variable. The standard deviation for both of these isotopic systems are at least twice larger in this group than for other groups buried in the convent (**Fig. 4**). The δ ^15^N and δ ^13^C values in bones and teeth correlate, suggesting that the individuals exhibiting the higher δ^15^N ratios could include more marine fish than the others into their diet [27]. In spite of the isotopic variability seen in the group, most individuals have δ^34^S values grouping between 8 to 9‰ (14/21). Among the individuals exhibiting those δ^34^S and for which Sr and O isotope values are available, the region of origin seems to be non-Breton Western France. The individual 20794, who comes from an inland region, and is younger than other individuals, is also clearly isotopically distinct from the others (**Fig. 4**). The other individuals coming from regions showing a similar δ^34^S values have a diet much poorer in animal proteins. Among the individuals with those δ^34^S values for which ^87^Sr/^86^Sr and δ^18^O values have been obtained, the geographical origin seems to have been the Alps or the Pyreneans. The δ^15^N of the mass Gr. 337 cannot be distinguished from that of the other local groups given the large range exhibited by those individuals. Overall, this high heterogeneity in isotopic composition suggests different diet, social status, and two main geographical origins: non-Breton Western France and French mountains or Switzerland.

## Discussion

### A small group of Breton soldiers

When using single, dual and triple isotope geographic assignments, 3 out of the 4 individuals from Gr. 322 (20193, 20185 and 20188) show a high probability of origin in Brittany (**Fig. 4**). However, we note that while the δ^34^S values in bone collagen from these same individuals are high, they are not identical to that of the teeth collagen. This observation suggests some regional mobility over the last decade of life within a mostly coastal region. The dual *δ*^18^O - ^87^Sr/^86^Sr assignment for the fourth individual (20183) excludes Brittany (**Fig. 6**). A more mainland origin of this individual is reinforced by the very low δ^34^S values in his tooth collagen (**Fig. 4**). Nevertheless, the much higher δ^34^S values in bone collagen also indicates that this individual moved to a more coastal region in its adult life (**Fig. 4**). Based on the combined isotopic data, the most likely region of origin of the childhood for this individual is North Central France, Southern France or central Spain (**Fig. 6**). This individual also shows the same specific mitochondrial haplogroup (H3+152) with two specific mutations (7805 and 16249) as two other individuals of Gr. 322 (20188, 20193) [16]. These specific polymorphisms are also found in the DNA of Louise de Quengo, a Breton aristocrat buried in the same convent 165 years later and who is also the local woman analyzed in our associated study [1, 56]. This shared mitochondrial haplogroup strongly suggests family ties within Brittany. Based on *δ*^15^N and *δ*^13^C data from teeth and bone collagen, two individuals from Gr. 322 (20188 and 20183), had a diet rich in animal protein suggesting a higher social status and, perhaps, a link to nobility (**Fig. 4**). They are also the tallest of the series with a mean stature of 1.72 m (**SI Text section Physical anthropology data**; fig. S2 and table S2). The fourth individual from Gr. 322 (20185) has all the isotopic characteristics of a local individual, but his diet seems to indicate a much more modest origin. We conclude that individuals from Gr. 322 are from the Breton camp. At least one of these individuals (20183) covered a large distance over the last few years while others might have traveled more regionally to the area of conflict. The presence of multiple old healed stab wounds and the intermediate δ^34^S values of individual 20183, suggest a relatively recent (few years at most) movement of a professional soldier allied to the Breton camp. Its diet rich in animal protein and mitochondrial haplogroup makes us favor the hypothesis of a noble military over a mercenary sent by Brittany’s allies described in historical sources [45,46,100,101].

### A Royal Army Tomb

We could only recover teeth of 6 individuals in Gr. 337, and only 4 of those 6 individuals had enough preserved collagen for δ^34^S analyses. Individuals from Gr. 337 display a much broader variety of natal origin than those in Gr. 322 when using dual *δ*^18^O - ^87^Sr/^86^Sr and triple isotope assignments (**Fig. 6**). Two individuals show high probability of origin during their childhood in western France including some regions of Brittany (20787; 20788). Teeth collagen was not preserved for these individuals and we could not validate if their δ^34^S values were also compatible with Brittany (**Fig. 6**). Two individuals show high probabilities of origin during their childhood from the northern Paris basin, Poitou region or Rhone valley (20781, 20765). Besides regions from mainland France, these individuals also show high probabilities of origin in regions of Spain, Flanders and England which could correspond to the allies of the Duchess Anne (the mercenaries sent, respectively, by Ferdinand of Aragon, Maximilian of Austria and Henry the VII). However, the mercenaries sent by different countries probably fought separately [46] making it unlikely that they would be buried together with their enemies in a mass grave. Two individuals show high probabilities of origin in the Alps excluding almost all other regions of Europe (20764, 20801). One of those individuals have been identified as a rider (20801) and another rider shows a similar bone δ^34^S. Based on the fact that armies from different regions fought separately, we can assume that all individuals from the grave 337 belong to the same political entity. Since all the individuals of the grave 337 are compatible with a French origin, we argue that individuals of the Gr. 337 correspond to soldiers of the royal camp recruited from different regions of Western mainland France and the Alps, which fits with historical sources [101]. In particular, the existence of soldiers coming from Switzerland and the Dauphine region has been documented during the French-Breton war and strongly argue for Gr. 337 to represent the Royal camp [101].

To validate this hypothesis, we further analyzed δ^34^S values in bones of 21 individuals from Gr. 337 (table S6). 18/21 individuals are incompatible with local Brittany δ^34^S values (**Fig. 4**) further indicating that these individuals moved a short time prior to their death to Rennes. The individuals have δ^34^S values in their bones between 5.3 and 12.6‰, which are all compatible with regions corresponding to the French Kingdom territory of that time, according to the δ^34^S isoscape that we developed for this study (**Fig. 4**). While some of these individuals could also originate from international regions allied with the Breton Camp, only 3 of them have values compatible with a coastal (and therefore possibly Breton) origin. One of them was a rider (20781), whom triple isoscape demonstrates a non-Breton origin (**Fig. 6**). The three lowest δ^34^S from bone or tooth also belongs to riders (**Figs. 4, 5**). Interestingly, the δ^34^S values of the non-rider individuals fall within a tight range between 7 and 10‰ (**Fig. 4**). While these values do not fit with Brittany, they are typical of western France and suggest a preferential recruitment from these regions of France for this conflict, which fits with historical documents mentioning soldiers coming from Poitou and Normandy (**Fig. 7**) [101]. δ^13^C and δ^15^N values from teeth collagen indicate a variable diet for individuals of Gr. 337 compared to other individuals buried in the convent (**Fig. 4**) [74]. While some individuals appear to have had a diet comparable to that of the social elites buried in the adjoining convent, others have the lowest values of δ^15^N in the series, reflecting a diet mostly based on plant sources. This disparity of diet further reinforces the idea that this grave gathered soldiers recruited from a variety of social status and geographical regions from mainland France.

**Figure 7.**
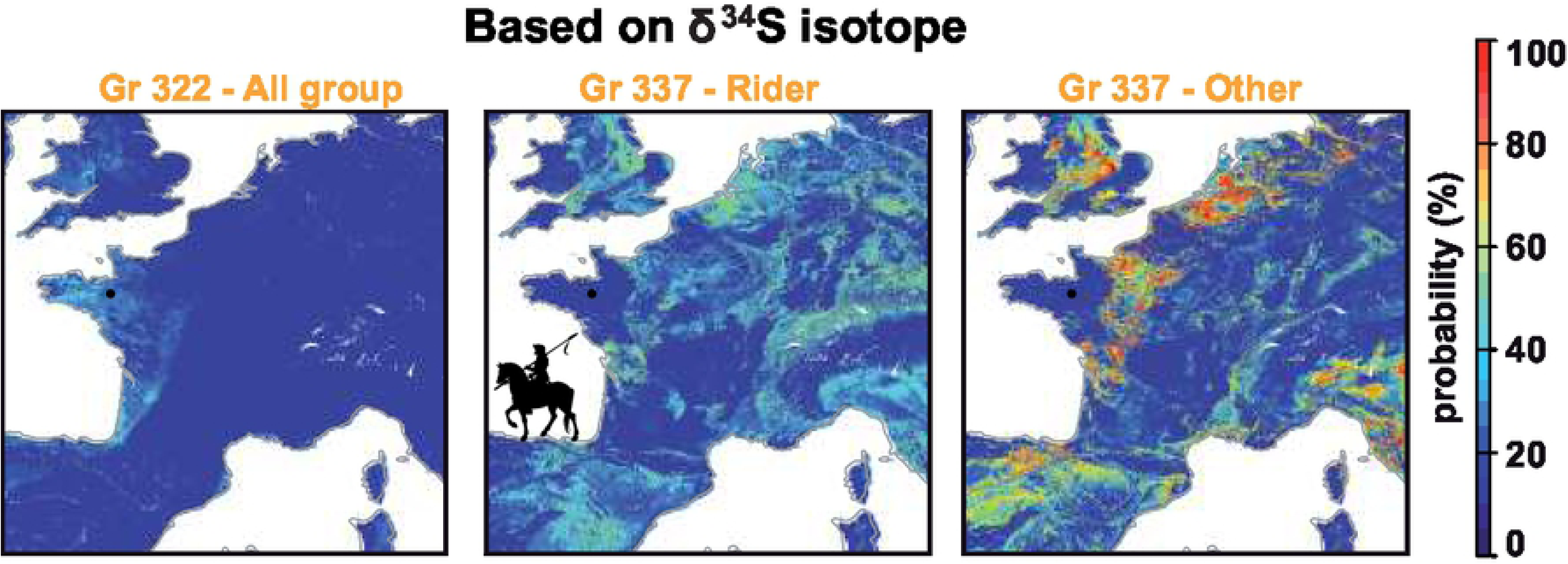
**Maps showing the location of residence of all recovered individuals (for which bones were available) towards the end of their life.** Maps of possible geographical origins are based on δ^34^S values. The probability maps are averaged by group: Gr. 322: 4 individuals; Gr 337 riders: 3 individuals identified as riders; Gr. 337; Other: 18 individuals. To generate these figures see **Bataille et al., in this issue** [1].

### Soldiers from different sides buried in the same place

These isotopic interpretations shed new light onto this undocumented conflict and raises new archeological questions. While other victims could have been buried elsewhere, an intriguing observation is the relative imbalance in the individual death count between the two graves, particularly when considering the defeat of the Brittany camp. The French soldiers (Gr. 337) are numerous, buried together and far from their home. Conversely, only four individuals were buried there from the Brittany camp. Based on the isotope interpretations, three of them are from regions distant from the battlefield. It is possible that only individuals with distant residences were buried on site explaining the higher Royal camp count. Another intriguing observation is the common burial place for soldiers of opposite camps. The high status and reputation of the Jacobins convent, also chosen to sign the peace treaty between King Charles VIII of France and Anne of Brittany on November 15, 1491 [42], may explain the choice of this place for these burials [50]. The presence of a famous and miraculous icon in the convent (“Notre-Dame de Bonne Nouvelle” meaning Our Lady of Good News) has undoubtedly guaranteed the sacred nature of the buildings and perhaps ensured an area of relative neutrality. The powerful Dominican friars were able to welcome in their cemetery the belligerents of both sides and allow, as suggested by the rosary and the absence of skinning, a slightly more careful inhumation than what is observed for most war victims of this time period [102]. Based on historical sources, the lethality of the conflict was limited [45].

Through a case study in Brittany, we demonstrated that a novel framework combining several traditional archeological tools and quantitative triple isotope geolocation techniques can shed light on unidentified and undocumented mass graves. Our interpretations provide critical insights into the origin, mobility and social status of buried individuals. This work, transferable to other graves, provides key tools for completing the missing historical archives of this conflict and resolving new archeological or forensic questions.

## Concluding remarks

Establishing human mobility using S isotopes is of high interest in archeology: this study shows that S isotopes can strongly complement other isotopic systems. It allows tracing migration that are often invisible with oxygen isotopes, which require large travel distances, and also further constraining provenance based on strontium isotopes, which can be redundant between distinct geographic regions. More than the simple identification of human provenance, our study shows that the use of this novel triple isotope geographic assignment in archeology can verify historical hypotheses on alliances and recruiting strategies in wars as well as improves historical archives by bringing information on the common soldiers’ life when historical documents usually focus on leaders. The combination of isotopes with DNA also reveal some surprises: we indeed report the existence of an individual with family ties in Brittany who grew up far from his region of origin, but traveled and lived in Brittany to fight the war threatening its independence. The isotopes geographic assignment is a real golden standard to complement archives on the mobility of unidentified human remains and to access hitherto previously unreleased information.

## Acknowledgments

Cost of isotope analyses was funded by the Max Planck Society. Salary support for Manuel Trost was provided by the Deutsche Forschungsgemeinschaft (DFG) PALEODIET (Project 378496604) and by the European Research Council (ERC) ARCHEIS (grant 803676) project for Klervia Jaouen. This work was also supported in part by National Sciences and Engineering Research Council (NSERC) Discovery Grant RGPIN-2019-05709 to Clement Bataille. We are grateful to Philip Miller and Hilary Bataille for editing the English of the manuscript.

## References

1. Bataille CP, Jaouen K, Milano S, Trost M, Steinbrenner S, Crubézy É, et al. Triple Sulfur-Oxygen-Strontium Isotopes Probabilistic Geographic Assignment of Archaeological Remains using a Novel Sulfur Isoscape of Western Europe. PLoS ONE. 2021.

2. Blanchard P, Castex D. A mass grave from the catacomb of Saints Peter and Marcellinus in Rome, second-third century AD. Antiquity. 2007; 989–998.

3. Duncan WN, Schwarz KR. A Postclassic Maya mass grave from Zacpetén, Guatemala. Journal of Field Archaeology. 2015;40: 143–165. doi:10.1179/0093469015Z.000000000113

4. McCormick M. Tracking mass death during the fall of Rome’s empire (I). Journal of Roman Archaeology. 2015;28: 325–357. doi:10.1017/S1047759415002512

5. Fiorato V, Boylston A, Knusel C, Erolin C. Blood red roses: The archaeology of a mass grave from the battle of Towton AD 1461. second. Oxford: Oxbow books; 2007. Available: https://discovery.dundee.ac.uk/en/publications/blood-red-roses-the-archaeology-of-a-mass-grave-from-the-battle-o

6. Castex D. Identification and Interpretation of Historical Cemeteries Linked to Epidemics. In: Raoult D, Drancourt M, editors. Paleomicrobiology: Past Human Infections. Berlin, Heidelberg: Springer Berlin Heidelberg; 2008. pp. 23–48. doi:10.1007/978-3-540-75855-6_2

7. Holst MR, Sutherland TL. Towton Revisited – Analysis of the Human Remains from the Battle of Towton 1461. In: Eickhoff S, Schopper F, editors. Schlachtfeld und Massengrab: Spektren Interdisziplinärer Auswertung von Orten der Gewalt. Zossen; 2014. pp. 97–129.

8. Meyer C, Lohr C, Gronenborn D, Alt KW. The massacre mass grave of Schöneck-Kilianstädten reveals new insights into collective violence in Early Neolithic Central Europe. PNAS. 2015;112: 11217–11222. doi:10.1073/pnas.1504365112

9. Biagini P, Thèves C, Balaresque P, Géraut A, Cannet C, Keyser C, et al. Variola Virus in a 300-Year-Old Siberian Mummy. New England Journal of Medicine. 2012;367: 2057–2059. doi:10.1056/NEJMc1208124

10. Thèves C, Cabot E, Bouakaze C, Chevet P, Crubézy É, Balaresque P. About 42% of 154 remains from the “Battle of Le Mans”, France (1793) belong to women and children: Morphological and genetic evidence. Forensic Sci Int. 2016;262: 30–36. doi:10.1016/j.forsciint.2016.02.029

11. Schroeder H, Margaryan A, Szmyt M, Theulot B, Włodarczak P, Rasmussen S, et al. Unraveling ancestry, kinship, and violence in a Late Neolithic mass grave. PNAS. 2019;116: 10705–10710. doi:10.1073/pnas.1820210116

12. Haak W, Brandt G, Jong HN de, Meyer C, Ganslmeier R, Heyd V, et al. Ancient DNA, Strontium isotopes, and osteological analyses shed light on social and kinship organization of the Later Stone Age. PNAS. 2008;105: 18226–18231. doi:10.1073/pnas.0807592105

13. Price TD, Johnson CM, Ezzo JA, Ericson J, Burton JH. Residential Mobility in the Prehistoric Southwest United States: A Preliminary Study using Strontium Isotope Analysis. Journal of Archaeological Science. 1994;21: 315–330. doi:10.1006/jasc.1994.1031

14. Bataille CP, Brennan SR, Hartmann J, Moosdorf N, Wooller MJ, Bowen GJ. A geostatistical framework for predicting variations in strontium concentrations and isotope ratios in Alaskan rivers. Chemical Geology. 2014;389: 1–15. doi:10.1016/j.chemgeo.2014.08.030

15. Willmes M, Bataille CP, James HF, Moffat I, McMorrow L, Kinsley L, et al. Mapping of bioavailable strontium isotope ratios in France for archaeological provenance studies. Applied Geochemistry. 2018;90: 75–86. doi:10.1016/j.apgeochem.2017.12.025

16. Thèves C, Mata X, Fromentier. Etude paléogénétique. Le Cloirec G. L’étude archéologique du couvent des Jacobins de Rennes (35), du quartier antique à l’établissement dominicain. Le Cloirec G. Cesson-Sévigné:; 2016. pp. 2219–2224.

17. Schwarcz HP, Gibbs L, Knyf M. Oxygen isotope analysis as an indicator of place of origin. In: Pfeiffer S, Williamson RE, editors. Snake Hill: An Investigation of a Military Cemetery from the War of 1812. Toronto: Dundurn Press; 1991. pp. 263–268.

18. White CD, Spence MW, Longstaffe FJ, Law KR. Testing the Nature of Teotihuacán Imperialism at Kaminaljuyú Using Phosphate Oxygen-Isotope Ratios. Journal of Anthropological Research. 2000;56: 535–558. doi:10.1086/jar.56.4.3630930

19. Dupras TL, Schwarcz HP. Strangers in a Strange Land: Stable Isotope Evidence for Human Migration in the Dakhleh Oasis, Egypt. Journal of Archaeological Science. 2001;28: 1199– 1208. doi:10.1006/jasc.2001.0640

20. White CD, Spence MW, Longstaffe FJ, Stuart-Williams H, Law KR. Geographic Identities of the Sacrificial Victims from the Feathered Serpent Pyramid, Teotihuacan: Implications for the Nature of State Power. Latin American Antiquity. 2002;13: 217–236. doi:10.2307/971915

21. White CD, Spence MW, Longstaffe FJ, Law KR. Demography and ethnic continuity in the Tlailotlacan enclave of Teotihuacan: the evidence from stable oxygen isotopes. Journal of Anthropological Archaeology. 2004;23: 385–403. doi:10.1016/j.jaa.2004.08.002

22. Chenery C, Müldner G, Evans J, Eckardt H, Lewis M. Strontium and stable isotope evidence for diet and mobility in Roman Gloucester, UK. Journal of Archaeological Science. 2010;37: 150–163. doi:10.1016/j.jas.2009.09.025

23. Evans JA, Chenery CA, Montgomery J. A summary of strontium and oxygen isotope variation in archaeological human tooth enamel excavated from Britain. Journal of Analytical Atomic Spectrometry. 2012;27: 754–764. doi:10.1039/C2JA10362A

24. King CL, Bentley RA, Tayles N, Viðarsdóttir US, Nowell G, Macpherson CG. Moving peoples, changing diets: isotopic differences highlight migration and subsistence changes in the Upper Mun River Valley, Thailand. Journal of Archaeological Science. 2013;40: 1681– 1688. doi:10.1016/j.jas.2012.11.013

25. Lightfoot E, O’Connell TC. On the Use of Biomineral Oxygen Isotope Data to Identify Human Migrants in the Archaeological Record: Intra-Sample Variation, Statistical Methods and Geographical Considerations. PLoS ONE. 2016;11: e0153850. doi:10.1371/journal.pone.0153850

26. Jaouen K, Colleter R, Pietrzak A, Pons M-L, Clavel B, Telmon N, et al. Tracing intensive fish and meat consumption using Zn isotope ratios: evidence from a historical Breton population (Rennes, France). Scientific reports. 2018;8: 5077.

27. Schoeninger MJ, Moore K. Bone stable isotope studies in archaeology. Journal of World Prehistory. 1992;6: 247–296. doi:10.1007/BF00975551

28. Buzon MR, Simonetti A, Creaser RA. Migration in the Nile Valley during the New Kingdom period: a preliminary strontium isotope study. Journal of Archaeological Science. 2007;34: 1391–1401. doi:10.1016/j.jas.2006.10.029

29. Knudson KJ, Price TD. Utility of multiple chemical techniques in archaeological residential mobility studies: Case studies from Tiwanaku- and Chiribaya-affiliated sites in the Andes. American Journal of Physical Anthropology. 2007;132: 25–39. doi:10.1002/ajpa.20480

30. Frei KM, Skals I, Gleba M, Lyngstrøm H. The Huldremose Iron Age textiles, Denmark: an attempt to define their provenance applying the strontium isotope system. Journal of Archaeological Science. 2009;36: 1965–1971. doi:10.1016/j.jas.2009.05.007

31. Eckardt H, Chenery C, Booth P, Evans JA, Lamb A, Müldner G. Oxygen and strontium isotope evidence for mobility in Roman Winchester. Journal of Archaeological Science. 2009;36: 2816–2825. doi:10.1016/j.jas.2009.09.010

32. Frei KM, Price TD. Strontium isotopes and human mobility in prehistoric Denmark. Archaeological and Anthropological Sciences. 2012;4: 103–114. doi:10.1007/s12520-011-0087-7

33. Scheeres M, Knipper C, Hauschild M, Schönfelder M, Siebel W, Pare C, et al. “Celtic migrations”: Fact or fiction? Strontium and oxygen isotope analysis of the Czech cemeteries of Radovesice and Kutná Hora in Bohemia. American Journal of Physical Anthropology. 2014;155: 496–512. doi:https://doi.org/10.1002/ajpa.22597

34. Gregoricka LA, Sheridan SG. Continuity or conquest? A multi-isotope approach to investigating identity in the Early Iron Age of the Southern Levant. American Journal of Physical Anthropology. 2016;162: 73–89. doi:https://doi.org/10.1002/ajpa.23086

35. Killgrove K, Montgomery J. All Roads Lead to Rome: Exploring Human Migration to the Eternal City through Biochemistry of Skeletons from Two Imperial-Era Cemeteries (1st-3rd c AD). PLOS ONE. 2016;11: e0147585. doi:10.1371/journal.pone.0147585

36. Knipper C, Mittnik A, Massy K, Kociumaka C, Kucukkalipci I, Maus M, et al. Female exogamy and gene pool diversification at the transition from the Final Neolithic to the Early Bronze Age in central Europe. PNAS. 2017 [cited 10 Nov 2020]. doi:10.1073/pnas.1706355114

37. Laffoon JE, Sonnemann TF, Shafie T, Hofman CL, Brandes U, Davies GR. Investigating human geographic origins using dual-isotope (87Sr/86Sr, δ18O) assignment approaches. PLOS ONE. 2017;12: e0172562. doi:10.1371/journal.pone.0172562

38. Scaffidi BK, Knudson KJ. An archaeological strontium isoscape for the prehistoric Andes: Understanding population mobility through a geostatistical meta-analysis of archaeological 87Sr/86Sr values from humans, animals, and artifacts. Journal of Archaeological Science. 2020;117: 105121. doi:10.1016/j.jas.2020.105121

39. Cavazzuti C, Skeates R, Millard AR, Nowell G, Peterkin J, Brea MB, et al. Flows of people in villages and large centres in Bronze Age Italy through strontium and oxygen isotopes. PLOS ONE. 2019;14: e0209693. doi:10.1371/journal.pone.0209693

40. Bataille CP, Holstein ICC von, Laffoon JE, Willmes M, Liu X-M, Davies GR. A bioavailable strontium isoscape for Western Europe: A machine learning approach. PLOS ONE. 2018;13: e0197386. doi:10.1371/journal.pone.0197386

41. Pichot D. Le temps des Montfort : « l’État breton ». Apogée. Bretagne est univers Catalogue du Musée de Bretagne. Apogée. Presses universitaires de Rennes; 2006. pp. 86–94.

42. Le Cloirec G. L’étude archéologique du couvent des Jacobins de Rennes (35), du quartier antique à l’établissement dominicain. Cesson-Sévigné: INRAP Grand-Ouest; 2016 p. 3835. Report No.: 12 vol.

43. Meyer J. Histoire de Rennes. 2e ed. Toulouse: Privat; 1984.

44. Leguay J-P, Martin H. Fastes et malheurs de la Bretagne ducale 1213-1532. Ouest-France; 1982.

45. Le Page D. 11 batailles qui ont fait la Bretagne. Morlaix: Skol Vreizh; 2015. Available: https://www.skolvreizh.com/catalog?page=shop.product_details&product_id=%24product_id

46. Currin JM. “The King’s Army into the Partes of Bretaigne”: Henry VII and the Breton Wars, 1489–1491. War in History. 2000;7: 379–412.

47. Albanao J, Cotton E. La dynastie des Habsbourg: La Maison d Autriche. Paris: Editions du Panthéon; 2012.

48. Charles VIII (King of France), La Trémoille LI de. Correspondance de Charles VIII et de ses conseillers avec Louis II de la Trémoille pendant la guerre de Bretagne, 1488. Genève: Mégariotis Reprints; 1978. Available: https://gallica.bnf.fr/ark:/12148/bpt6k7065m

49. Calmette J. La politique espagnole dans la crise de l’indépendance bretonne (1488-1492). Revue Historique. 1914;117: 168–182.

50. Martin H. Les ordres mendiants en Bretagne (vers 1230 - vers 1530). Pauvreté volontaire et prédication à la fin du Moyen-Âge. Rennes: Librairie C. Klincksieck, 11 rue de Lille, Paris; 1975.

51. Mokrane FZ, Colleter R, Duchesne S, Gerard P, Savall F, Crubézy É, et al. Old hearts for modern investigations: CT and MR for archaeological human hearts remains. Forensic Science International. 2016; 14–24. doi:10.1016/j.forsciint.2016.08.035

52. Vovelle M. La mort et l’Occident. De 1300 à nos jours. Paris: Gallimard; 2000.

53. Boissavit-Camus B, Zadora-Rio E. L’organisation spatiale des cimetières paroissiaux. In: Galinié H, Zadora-Rio E, editors. Archéologie du cimetière chrétien - Actes du 2ème colloque ARCHEA Orléans, 29 septembre-1er octobre 1994. Tours: 11e supplément à la Revue Archéologique de Centre de la France - Conseil Régional du Centre; 1996. pp. 49– 53.

54. Ariès P. Hour of Our Death. Vintage Books; 1981.

55. Colleter R. Pratiques funéraires, squelettes et inégalités sociales. Etude d’un échantillon des élites bretonnes à l’époque moderne. phd, Université de Toulouse, Université Toulouse III - Paul Sabatier. 2018. Available: http://thesesups.ups-tlse.fr/4159/

56. Colleter R, Dedouit F, Duchesne S, Mokrane F-Z, Gendrot V, Gérard P, et al. Procedures and Frequencies of Embalming and Heart Extractions in Modern Period in Brittany. Contribution to the Evolution of Ritual Funerary in Europe. PLoS ONE. 2016;11: e0167988. doi:10.1371/journal.pone.0167988

57. Colleter R, Dedouit F, Duchesne S, Gérard P, Dercle L, Poilpré P, et al. Study of a seventeenth-century French artificial mummy: autopsical, native, and contrast-injected CT investigations. International Journal of Legal Medicine. 2018;132: 1405–1413. doi:10.1007/s00414-018-1830-8

58. Bruzek J. A Method for Visual Determination of Sex, Using the Human Hip Bone. American Journal of Physical Anthropology. 2002;117: 157–168. doi:10.1002/ajpa.10012.abs

59. Murail P, Bruzek J, Houët F, Cunha E. DSP: a tool for probabilistic sex diagnosis using worldwide variability in hip-bone measurements. Bulletins et Mémoires de la Société d’Anthropologie de Paris. 2005; 167–176.

60. Schmitt A. Une nouvelle méthode pour estimer l’âge au décès des adultes à partir de la surface sacro-pelvienne iliaque. Bulletins et Mémoires de la Société d’Anthropologie de Paris. 2005;17: 89–101.

61. Moorrees CFA, Fanning EA, Hunt EE. Formation and resorption of three deciduous teeth in children. American Journal of Physical Anthropology. 1963;21: 205–213. doi:10.1002/ajpa.1330210212

62. Moorrees CFA, Fanning EA, Hunt EE. Age variation of formation stages for ten permanent teeth. Journal of dental Research. 1963;42: 1490–1502.

63. Birkner R. L’image radiologique typique du squelette : aspect normal et variantes chez l’adulte et l’enfant ; pour médecins, étudiants et manipulateurs. Paris: Maloine; 1980.

64. Owings Webb PA, Myers Suchey J. Epiphyseal union of the anterior iliac cret and medial clavicule in a modern multiracial sample of american males and females. American Journal of Physical Anthropology. 1985;68: 457–466.

65. Ruff CB, Holt BM, Niskanen M, Sladék V, Berner M, Garofalo E, et al. Stature and body mass estimation from skeletal remains in the European Holocene. American Journal of Physical Anthropology. 2012;148: 601–617. doi:10.1002/ajpa.22087

66. Duday H, Courtaud P, Crubézy É, Sellier P, Tillier A-M. L’Anthropologie « de terrain » : reconnaissance et interprétation des gestes funéraires. Bulletins et Mémoires de la Société d’anthropologie de Paris. 1990;2: 29–49. doi:10.3406/bmsap.1990.1740

67. Aufderheide AC, Rodriguez-Martin C. The Cambridge Encyclopedia of Human Paleopathology. Cambridge University Press; 1998.

68. Capasso B. Brucellosis at Herculanum (79 AD). International journal of osteoarcheology. 1999;9: 277–288.

69. Ortner DJ, editor. Identification of Pathological Conditions in Human Skeletal Remains. Second. San Diego CA: Academic Press; 2003. Available: http://www.sciencedirect.com/science/article/pii/B9780125286282500612

70. Pinhasi R, Mays S, editors. Advances in Human Palaeopathology. Chichester: John Wiley & Sons; 2008. Available: https://books.google.fr/books/about/Advances_in_Human_Palaeopathology.html?hl=fr&id=N_hVqemQCNUC

71. Waldron T. Palaeopathology. Cambridge: Cambridge University Press; 2009.

72. Villotte S. Enthésopathies et activités des hommes préhistoriques : recherche méthodologique et application aux fossiles européens du Paléolithique supérieur et du Mésolithique. Oxford: BAR International Series; 2009. Available: https://books.google.fr/books/about/Enth%C3%A9sopathies_et_activit%C3%A9s_des_hommes.html?hl=fr&id=D7USQgAACAAJ

73. Capuani C, Telmon N, Moscovici J, Molinier F, Aymeric A, Delisle M-B, et al. Modeling and determination of directionality of the kerf in epifluorescence sharp bone trauma analysis. Int J Legal Med. 2014;128: 1059–1066. doi:10.1007/s00414-014-1022-0

74. Colleter R, Clavel B, Pietrzak A, Duchesne S, Schmitt L, Richards MP, et al. Social status in late medieval and early modern Brittany: insights from stable isotope analysis. Archaeological and Anthropological Sciences. 2019;11: 823–837.

75. Talamo S, Richards M. A Comparison of Bone Pretreatment Methods for AMS Dating of Samples \textgreater30,000 BP. Radiocarbon. 2011;53: 443–449.

76. Deniel C, Pin C. Single-stage method for the simultaneous isolation of lead and strontium from silicate samples for isotopic measurements. Analytica Chimica Acta. 2001;426: 95– 103.

77. Leclerc J. La notion de sépulture. Bulletins et Mémoires de la Société d’Anthropologie de Paris. 1990;2: 13–18. doi:10.3406/bmsap.1990.1738

78. Knüsel CJ, Robb J. Funerary taphonomy: An overview of goals and methods. Journal of Archaeological Science: Reports. 2016;10: 655–673. doi:10.1016/j.jasrep.2016.05.031

79. Vanezis P. Investigation of clandestine graves resulting from human rights abuses. Journal of Clinical Forensic Medicine. 1999;6: 238–242. doi:10.1016/S1353-1131(99)90004-4

80. Komar D. Differential decay rates in single, multiple, and mass graves in Bosnia. Proceedings of the American Academy of Forensic Sciences Annual Meeting,. 2001;7: 242– 243.

81. Jessee E, Skinner M. A typology of mass grave and mass grave-related sites. Forensic Science International. 2005;152: 55–59. doi:10.1016/j.forsciint.2005.02.031

82. Dutour O, Jankauskas R, Buzhilova AP. Ensembles funéraires liés à des faits de guerre. Cimetières et identités. Bordeaux: Ausonius; 2015. pp. 133–146. Available: https://dokupdf.com/download/est-paru-cimetieres-et-identites-_5a01b514d64ab2b9bd6733e5_pdf

83. Nicklisch N, Ramsthaler F, Meller H, Friederich S, Alt KW. The face of war: Trauma analysis of a mass grave from the Battle of Lützen (1632). PLoS ONE. 2017;12: e0178252. doi:10.1371/journal.pone.0178252

84. Walker PL. A Bioarchaeological Perspective on the History of Violence. Annual Review of Anthropology. 2001;30: 573–596. doi:10.1146/annurev.anthro.30.1.573

85. Murphy K, Waa S, Jaffer H, Sauter A, Chan A. A literature review of findings in physical elder abuse. Canadian Association of Radiologists Journal = Journal l’Association Canadienne Des Radiologistes. 2013;64: 10–14. doi:10.1016/j.carj.2012.12.001

86. Flieger A, Kölzer SC, Plenzig S, Heinbuch S, Kettner M, Ramsthaler F, et al. Bony injuries in homicide cases (1994-2014). A retrospective study. International Journal of Legal Medicine. 2016;130: 1401–1408. doi:10.1007/s00414-016-1407-3

87. Schreider E. Consanguinité et variations biologiques chez l’homme. Population. 1976;31: 341–354. doi:10.2307/1530447

88. Marquer P. Endogamie, exogamie et variations de la stature et de l’indice céphalique dans la population béarnaise (Pyrénées-Atlantiques). Bulletins et Mémoires de la Société d’Anthropologie de Paris. 1979;6: 333–342. doi:10.3406/bmsap.1979.1969

89. Olivier G, Devigne G. Consanguinity and endogamy. Journal of Human Evolution. 1980;9: 261–268. doi:10.1016/0047-2484(80)90054-8

90. Susanne C. Living conditions and secular trend. Journal of Human Evolution. 1985;14: 357– 370. doi:10.1016/S0047-2484(85)80042-7

91. Vercauteren M. Evolution séculaire au XXe siècle. Anthropologie biologique Evolution et biologie humaine. De Boeck Université; 2003. pp. 539–547.

92. Heyberger L. Stature et niveau de vie biologique des conscrits du Limousin (1782-1940). Un indice de développement socio-économique. Histoire & Sociétés Rurales. 2005;24: 83– 104.

93. Susanne C, Polet C. Dictionnaire d’anthropobiologie. De Boeck. Bruxelles: De Boeck Université; 2005.

94. Burgess A. The excavation and finds. In: Fiorato V, Boylston A, Knusel C, editors. Blood Red Roses: The Archaeology of a Mass Grave from the Battle of Towton, AD 1461. Oxford: Oxbow Books; 2000. pp. 29–35.

95. Hillson S. Dental anthropology. Cambridge: Cambridge University Press; 2008.

96. Hedges REM, Clement JG, Thomas CDL, O’Connell TC. Collagen turnover in the adult femoral mid-shaft: Modeled from anthropogenic radiocarbon tracer measurements. American Journal of Physical Anthropology. 2007;133: 808–816. doi:10.1002/ajpa.20598

97. Nehlich O, Borić D, Stefanović S, Richards MP. Sulphur isotope evidence for freshwater fish consumption: a case study from the Danube Gorges, SE Europe. Journal of Archaeological Science. 2010;37: 1131–1139. doi:10.1016/j.jas.2009.12.013

98. Nehlich O, Fuller BT, Jay M, Mora A, Nicholson RA, Smith CI, et al. Application of sulphur isotope ratios to examine weaning patterns and freshwater fish consumption in Roman Oxfordshire, UK. Geochimica et Cosmochimica Acta. 2011;75: 4963–4977. doi:10.1016/j.gca.2011.06.009

99. Privat KL, O’Connell TC, Hedges REM. The distinction between freshwater- and terrestrial-based diets: methodological concerns and archaeological applications of sulphur stable isotope analysis. Journal of Archaeological Science. 2007;34: 1197–1204. doi:10.1016/j.jas.2006.10.008

100. Jones M. L’armée bretonne 1449-1491 : Structures et carrières. La France de la fin du xve siècle : renouveau et apogée. Paris: CNRS Editions; 1985. pp. 147–166.

101. Contamine P. Chapitre X. L’armée du Roi de France : vue d’ensemble. Guerre, État et société à la fin du Moyen Âge Tome 1 : Études sur les armées des rois de France 1337-1494. Paris: Éditions de l’École des hautes études en sciences sociales; 2013. pp. 277–319. Available: http://books.openedition.org/editionsehess/700

102. Komar D. Patterns of Mortuary Practice Associated with Genocide: Implications for Archaeological Research. Current Anthropology. 2008;49: 123–133. doi:10.1086/524761

